# Rational Design of 5’ UTRs for Optimized mRNA Expression and Cost-Effective, Rapid Analysis of Capped mRNA

**DOI:** 10.64898/2026.10.08.757771

**Authors:** Yuan Fang, Tao Liu, Yiwen Liang, Xiaodie Wang, Meiqi Shao, Congcong Wang, Jianchen Yu, Yan Sun, Pingjing Zhang, Qijun Qian

## Abstract

In vitro-transcribed (IVT) messenger RNA (mRNA) therapeutics require rigorous quality control of 5’ capping efficiency and 3’ poly(A) tail length, both of which constitute critical quality attributes (CQAs). Current analytical workflows for these CQAs rely on hybridization with nucleic acid probes and biotin-based affinity purification, which constrain throughput and increase per-assay cost. Here, we report a rationally designed 5’ untranslated region (5’ UTR) that sustains high translational efficiency while enabling probe-free, one-step purification for rapid quantification of capping efficiency and poly(A) tail length distribution. This approach exploits a strategically engineered 5’ UTR incorporating a purine-only RNase A-resistant fragment terminating at a single pyrimidine cleavage site, together with a translation-enhancing element comprising an N^6^-methyladenosine (m^6^A) deposition motif and an 18S rRNA complementarity sequence. This dual-function design accommodates GA-rich sequence segments without compromising mRNA translational output. Compared with conventional probe- and biotin-dependent workflows, the present method substantially reduces cost and processing time and is compatible with simultaneous detection of homopolymeric poly(A) tail. This integrated platform provides a simplified, probe-free and construct-universal mRNA analytical platform.

## Introduction

Messenger RNA (mRNA)-based vaccines and therapeutics have emerged as a transformative modality in precision medicine, enabling expression of intracellular, transmembrane, and secreted proteins without the risk of genomic integration^1–10^. The rapid development and global deployment of mRNA-1273 (Moderna) and BNT162b2 (Pfizer–BioNTech), with billions of doses administered, underscore the agility and scalability of this platform^11–13^. As mRNA candidates advance through clinical pipelines, rigorous characterization of critical quality attributes (CQAs), particularly 5’ capping efficiency and 3’ poly(A) tail length distribution, has become essential for regulatory compliance and manufacturing scale-up^14–16^. Accordingly, the development of accurate, robust, and operationally streamlined analytical methods for quantifying capping efficiency and poly(A) tail length distribution is indispensable for process development, batch release, and regulatory compliance of mRNA drug substances.

The 5’ cap structure is essential for mRNA function, recruiting eukaryotic initiation factor 4E (eIF4E) for translation initiation and conferring resistance to 5’→3’ exonucleolytic degradation. Current methods for determining capping efficiency each carry intrinsic limitations. ^32^P-radiolabeling, although highly sensitive, requires radioactively labeled mRNA and is incompatible with current Good Manufacturing Practice (cGMP) environments^17^. Enzyme-linked immunosorbent assay (ELISA)-based approaches using anti-m⁷G antibodies are operationally straightforward but cannot discriminate among GpppN (uncapped guanosine), Cap0 (m⁷GpppN), and Cap1 (m⁷GpppNm) structures (Fig.1), as the antibody recognizes the common m^7^G moiety common to both capped species and does not detect uncapped GpppN termini^15,18^. Nuclease digestion coupled with reverse-phase high-performance liquid chromatography (RP-HPLC) can separate capped from uncapped fragments but lacks the resolving power to baseline-separate GpppN, Cap0, and Cap1, which differ by only a single methyl group^19,20^. The RNase H-based probe method is currently regarded as the benchmark for capping efficiency analysis. It employs a custom-designed chimeric DNA–RNA probe to site-specifically cleave the mRNA 5’ terminus, followed by streptavidin magnetic-bead enrichment and liquid chromatography-tandem mass spectrometry (LC–MS/MS) quantification of the released cap-bearing oligonucleotide^19,21^. Although this approach offers high cleavage specificity and quantitative accuracy, two practical constraints limit its utility in high-throughput settings: the DNA-containing probe must be custom-designed for each mRNA sequence, and probe synthesis typically requires 7–10 business days, imposing substantial lead-time and material costs^21^. Recent efforts have sought to address these limitations using ribozymes or DNAzymes for site-specific RNA cleavage, yet these catalysts require substrate-specific design and exhibit variable cleavage efficiency across different mRNA sequences^15,22,23^.

**Fig. 1.**
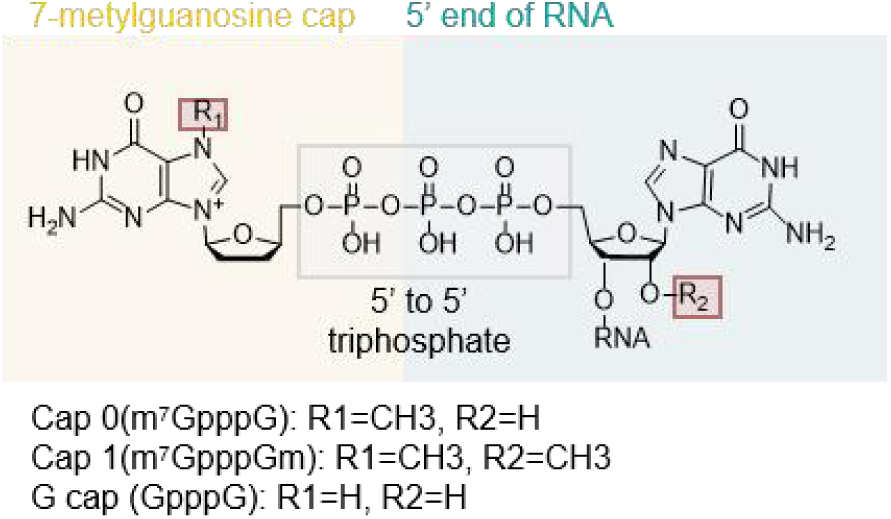
The 5’ terminal cap structure of mRNA. A guanylate (G) is added to the 5’ end of the mRNA to form G-CAP. On the basis of G-CAP, the 7th nitrogen atom of guanine is methylated (7-methylguanine, m⁷G) to form Cap0. Once a Cap 0 structure has been produced, it can be further modified by 2’-Oribose methyltransferase to result in a Cap 1 structure.

The 3’ poly(A) tail interacts with poly(A)-binding protein (PABP) to protect mRNA from 3’→5’ exonucleolytic decay^24–26^ and enhances translational efficiency through the closed-loop ribosomal recycling model^27^. For IVT mRNA, poly(A) tail lengths of 100–150 nt are generally considered optimal, as deviations in either direction correlate with reduced translational output and accelerated intracellular degradation. Existing methods for poly(A) tail characterization present significant analytical constraints. Size-exclusion chromatography (SEC) provides a weight-average tail length but yields no information on length distribution or 3’ end heterogeneity^14^. IP-RP-LC improves resolution of poly(A) tail variants up to approximately 150 nt but requires RNase T1 pre-digestion to liberate the tail fragment and cannot simultaneously assess 5’ capping efficiency^14^. Sequencing-based approaches, including TAIL-seq^28^, PAL-seq^29^, and direct RNA sequencing (DRS)^30^, offer nucleotide-level tail length profiles but require laborious library preparation, bioinformatic pipelines, and internal length calibration standards, rendering them impractical for routine QC. Furthermore, all of the above methods assess 5’ capping efficiency and 3’ poly(A) tail length as independent analytical workflows, requiring separate sample preparation, probe design, and instrument time.

Here, we describe a fundamentally different strategy that exploits the intrinsic specificity of RNase A, which cleaves RNA on the 3’ side of pyrimidine residues, to achieve simultaneous dual-CQA analysis without any sequence-specific regent. The approach is enabled by rationally engineering the mRNA 5’ untranslated region (5’ UTR) to incorporate a purine-only segment (Part A, 9–27 nt stretch composed exclusively of A and G residues) that is intrinsically resistant to RNase A, flanked by a single terminal pyrimidine that serves as the sole cleavage site. Concurrently, a translation-enhancing element comprising an N6-methyladenosine (m^6^A) deposition motif and an 18S ribosomal RNA (rRNA) complementarity sequence sustains high translational output despite the GA-rich Part A architecture. The 3’ poly(A) tail is engineered as a segmented AₘGₙ heteropolymeric structure that also resists RNase A digestion. Following a single-step RNase A digestion and silica spin-column purification, LC-MS/MS analysis resolves GpppN, Cap0, and Cap1 species by accurate mass and simultaneously determines poly(A) tail length from the intact product. Once Part A is incorporated into the mRNA template, no construct-specific reagent design, probe synthesis, or catalyst customization is required, and the same RNase A reagent is universally applicable across mRNA constructs. The same RNase A reagent is universally applicable across all mRNA constructs. This integrated assay eliminates the need for custom probe synthesis and magnetic-bead enrichment, substantially reduces per-sample processing time and cost, and provides a streamlined analytical platform.

## Materials and Methods

### Plasmid construction and RNA31 preparation

DNA oligonucleotides RNA31-F and RNA31-R (Supplementary Table S1, SEQ ID NO: 1 and 2, respectively) were commercially synthesized (General Bio, Anhui, China) and annealed to generate double-stranded RNA31.

For in vitro transcription templates, eGFP and Fluc reporter constructs were synthesized with either the mRNA-1273 (Moderna-derived reference, MDR) 5’ UTR (Supplementary Table S1, SEQ ID NO: 4 and 5) or synthetic 5’ UTR variants T1-T10 (Supplementary Table S1, SEQ ID NO: 6-15). These cassettes were cloned into a backbone vector containing a T7 promoter, the synthetic 3’-UTR 36H2U (Supplementary Table S1, SEQ ID NO: 16), and a poly(A) sequence containing a 62GG motif (Supplementary Table S1, SEQ ID NO: 17). The resulting plasmids were linearized with NheI prior to transcription. Transcribed and capped RNAs were designated MDR, vMDR, and T1-T10 (Supplementary Table S1, SEQ ID NO: 18-29). Detailed sequence information for these transcription templates is archived in Supplementary Table S1.

### In vitro transcription and enzymatic capping

IVT was performed using a commercial T7 RNA polymerase kit according to the manufacturer’s protocol. The 20 μL reaction contained 10× buffer, ATP, CTP, pseudo-UTP, GTP (2 μL each), T7 polymerase (1.5 μL), 1.5 mol/L MgCl_2_ (0.27 μL), and 5 μg NheI-linearized template encoding a homopolymeric poly(A) tract. After incubation at 37℃ for 4 h, residual DNA was digested with DNase I (5 U per μg template) at 37℃ for 30–60 min. The mRNA was purified by lithium chloride precipitation (7.5 mol/L LiCl, −20℃, 30 min; 5,000 rpm at 4℃ for 20 min), washed with 75% (v/v) ethanol, dried, and resuspended in nuclease-free water. For enzymatic capping, the mRNA was denatured at 65℃ for 5 min, chilled on ice, and incubated for 2 h with vaccinia capping enzyme (VCE, 1μL) mix, 32 mmol/L S-adenosylmethionine (SAM, 0.13μL), and 100 mmol/L GTP-Na_2_ (0.1μL), yielding predominantly Cap1 (m^7^GpppNmN) with minor Cap0 (m^7^GpppN). The capped product was repurified by LiCl precipitation for subsequent capping efficiency and poly(A) tail length analysis.

### Simultaneous determination of capping efficiency and poly(A) tail length by RNase A digestion and LC-MS analysis

mRNA samples (80 μg) were digested in a 100 μL reaction mixture containing 2.5 μL RNase A (Biolabs, Ipswich, MA, USA) and 33.3 μL 10× RNase A buffer (100 mmol/L Tris-HCl, pH 7.5; 3 mol/L NaCl; 50 mmol/L EDTA), diluted to 3× working concentration with nuclease-free water. The reaction was incubated at 37℃ for 1 h. The digestion products were purified using the Monarch^®^ RNA Cleanup Kit (New England Biolabs, Ipswich, MA, USA) according to the manufacturer’s protocol. The eluate was analyzed by ultra-performance liquid chromatography-quadrupole time-of-flight mass spectrometry (UPLC-Q-TOF-MS; Waters Corporation, Milford, MA, USA) on an ACQUITY UPLC Oligonucleotide BEH C18 column (130 Å, 1.7 μm, 2.1 × 100 mm) maintained at 65℃ with the autosampler at 10°C. Mobile phase A consisted of 0.75% hexafluoroisopropanol (HFIP), 0.0375% N,N-diisopropylethylamine (DIEA), and 10 μmol/L EDTA in water; mobile phase B was prepared as 20:80 (v/v) water/methanol with identical additives. Gradient elution was performed at 0.3 mL/min (5% B initially, held for 2 min; linear increase to 50% B at 10 min; step to 90% B at 10.5 min, held to 12.5 min; return to 5% B at 13 min, equilibrated to 15 min). UV detection was monitored at 260 nm. Mass spectrometric acquisition was conducted in negative-ion full-scan mode (m/z 400-3000) with a scan speed of 2 Hz, capillary voltage of 2.5 kV, cone voltage of 40 V, and desolvation temperature of 400℃. Capping efficiency was quantified by extracted ion chromatogram peak area normalization of cap species (triphosphate, monophosphate, diphosphate, Cap0, Cap1), whereas poly(A) tail length distribution was determined from the corresponding oligoadenylate ion series.

### Capping efficiency determination by RNase H-directed cleavage and UPLC-MS analysis

A 5’-biotinylated DNA oligonucleotide probe (Supplementary Table S1, SEQ ID NO: 50) complementary to the 5’-UTR of the target mRNA was designed and annealed to the mRNA at a 1:1 molar ratio. The mixture was denatured at 85℃ for 1 min and subjected to stepwise gradient cooling (65℃, 55℃, and 40℃, 2 min each) to 22℃ to form an RNA/DNA heteroduplex. The hybridized substrate was incubated with thermostable RNase H (5 U; New England Biolabs, Ipswich, MA, USA) at 37℃ for 2 h to selectively cleave the RNA strand at the 5’ terminus, liberating oligonucleotide fragments bearing distinct cap structures (triphosphate, monophosphate, diphosphate, Cap0, and Cap1). The biotinylated 5’ fragments were captured by incubation with streptavidin-coated magnetic beads (Dynabeads MyOne Streptavidin C1, 10 mg/mL; Invitrogen, Waltham, MA, USA) at ambient temperature for 30 min. The beads were washed twice with 2× B&W buffer and twice with nuclease-free water, followed by elution at 85℃ for 3 min in 1 mmol/L EDTA containing 1% (v/v) methanol. The eluate was centrifuged at 12,000 × g for 10 min, and the supernatant was analyzed by ultra-performance liquid chromatography–mass spectrometry (UPLC-MS; Waters Corporation, Milford, MA, USA) on an ACQUITY Premier Oligonucleotide BEH C18 column (2.1 × 100 mm, 1.7 μm). Chromatographic separation was achieved using mobile phase A (0.75% hexafluoroisopropanol (HFIP), 0.0375% N,N-diisopropylethylamine (DIEA), and 10 μmol/L EDTA in water) and mobile phase B (acetonitrile) under gradient elution (5–50% B over 0–10 min; 90% B at 10.5–12.5 min) at a flow rate of 0.3 mL/min and column temperature of 65℃. Mass spectrometric detection was performed in negative-ion full-scan mode (m/z 400–3000) with a capillary voltage of 2.5 kV and desolvation temperature of 400℃. Capping efficiency was calculated by peak area normalization of extracted ion chromatograms corresponding to each cap species.

Capping efficiency calculation:

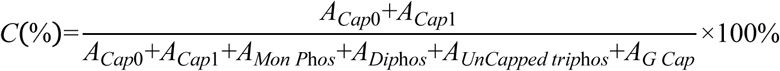

The relative abundance of each 5’-terminal species was quantified from the deconvoluted mass spectrum of the cap region. Extracted peak intensities were assigned as follows: *A_Cap_*_0_, intensity of Cap0 (m^7^GpppN); *A_Cap_*_1_, intensity of Cap1 (m⁷GpppNmN); *A_Mon P_*_h*os*_, intensity of 5’-monophosphate (pN); *A_Dip_*_h*os*_ , intensity of 5’-diphosphate (ppN); *A_UnCapped trip_*_h*os*_ , intensity of uncapped triphosphate (pppN); and *A_G Cap_* , intensity of unmethylated G cap (GpppN).

### Poly(A) tail length analysis by RNase A digestion and magnetic bead capture

The poly(A) tail was enriched and isolated from full-length mRNA via selective RNase A (New England Biolabs, Ipswich, MA, USA) digestion coupled with oligo(dT) magnetic bead (Nanomicro, Suzhou, China) capture. Briefly, oligo(dT) magnetic beads (10 mg/mL, 50 μL) were equilibrated with binding buffer (100 μL, twice) and resuspended in 80 μL binding buffer. The mRNA sample (approximately 50 pmol) was mixed with 5 M NaCl (20 μL) and nuclease-free water (Tiangen, CN) to a final volume of 80 μL, then incubated with the pre-treated beads under gentle agitation at ambient temperature for 10 min to facilitate hybridization of the poly(A) tail to oligo(dT). Following magnetic separation, RNase A (2 μL) was added to the bead-bound mRNA complex and incubated at 37℃ for 1 h to selectively digest single-stranded regions containing cytidine and uridine residues, thereby preserving the poly(A) tract. The beads were sequentially washed with binding buffer (100 μL, two cycles) and 100 mM ammonium acetate (100 μL, two cycles) to remove digested fragments and residual enzyme. The enriched poly(A) tail was eluted at 85℃ for 3 min in 30 μL of 100 μmol/L EDTA containing 1% (v/v) methanol in nuclease-free water, followed by magnetic separation. The eluate was transferred to a fresh microcentrifuge tube and centrifuged at 12,000 × g for 10 min at ambient temperature. The supernatant was collected for downstream analysis by liquid chromatography–tandem mass spectrometry (LC-MS/MS; Waters Corporation, Milford, MA, USA). The elution volume may be adjusted according to experimental requirements.

### Reporter-based assessment of mRNA translation efficiency

Human embryonic kidney 293T (HEK293T) cells (ATCC, USA) were cultured in Dulbecco’s Modified Eagle Medium (DMEM; Corning, USA) supplemented with 10% (v/v) fetal bovine serum (FBS; Gibco, USA) and 1% (v/v) penicillin-streptomycin (Thermo Fisher Scientific, USA) at 37℃ in a humidified atmosphere containing 5% CO₂. For reporter gene assays, cells at logarithmic growth phase were aliquoted at 1 × 10^6^ cells per sample. eGFP- or Fluc-encoding mRNA (5 μg) was transfected using the SF Cell Line 4D-Nucleofector X Kit (V4XC-2032; Lonza, Switzerland) with program CM-130. At day 2, day 4, and day 6 post-electroporation, cells were washed with phosphate-buffered saline (PBS), detached with 0.5% (w/v) trypsin-EDTA, and harvested. eGFP fluorescence was quantified by flow cytometry as mean fluorescence intensity (MFI) and positive percentage, whereas Fluc activity was determined using D-luciferin potassium salt to assess mRNA translation efficiency.

### Statistical analysis

Statistical analyses were carried out using GraphPad Prism 10.0 (GraphPad Software, San Diego, CA). Data were analyzed using unpaired *t* tests to compare two different conditions and analysis of variance for more conditions. All error bars show the standard errors of the means (SEM). Statistics significant difference was considered as following: P≥ 0.05 (ns), P < 0.05 (*), P < 0.01 (**), P < 0.001 (***), P<0.0001 (****).

## Results

### RNase A digestion preserves the 5’ cap structure of mRNA

RNase A is a pancreatic endoribonuclease that catalyzes transphosphorylation of the 3’, 5’-phosphodiester bond exclusively at the 3’ side of pyrimidine residues (C and U) in single-stranded RNA^31^. Under high-salt conditions, RNase A selectively hydrolyzes single-stranded RNA while sparing double-stranded regions and RNA-DNA heteroduplexes^32,33^. This strict pyrimidine 3’-specificity provides a predictable biochemical logic: any RNA segment devoid of C and U residues is intrinsically resistant to RNase A, regardless of its sequence context or reaction conditions. We leveraged this property to engineer an mRNA architecture in which the 5’ cap-bearing region and the poly(A) tail are both encoded within RNase A-resistant, purine-only segments, while all intervening sequences are designed to contain regular pyrimidine cleavage sites (Fig. 2A).

**Fig. 2.**
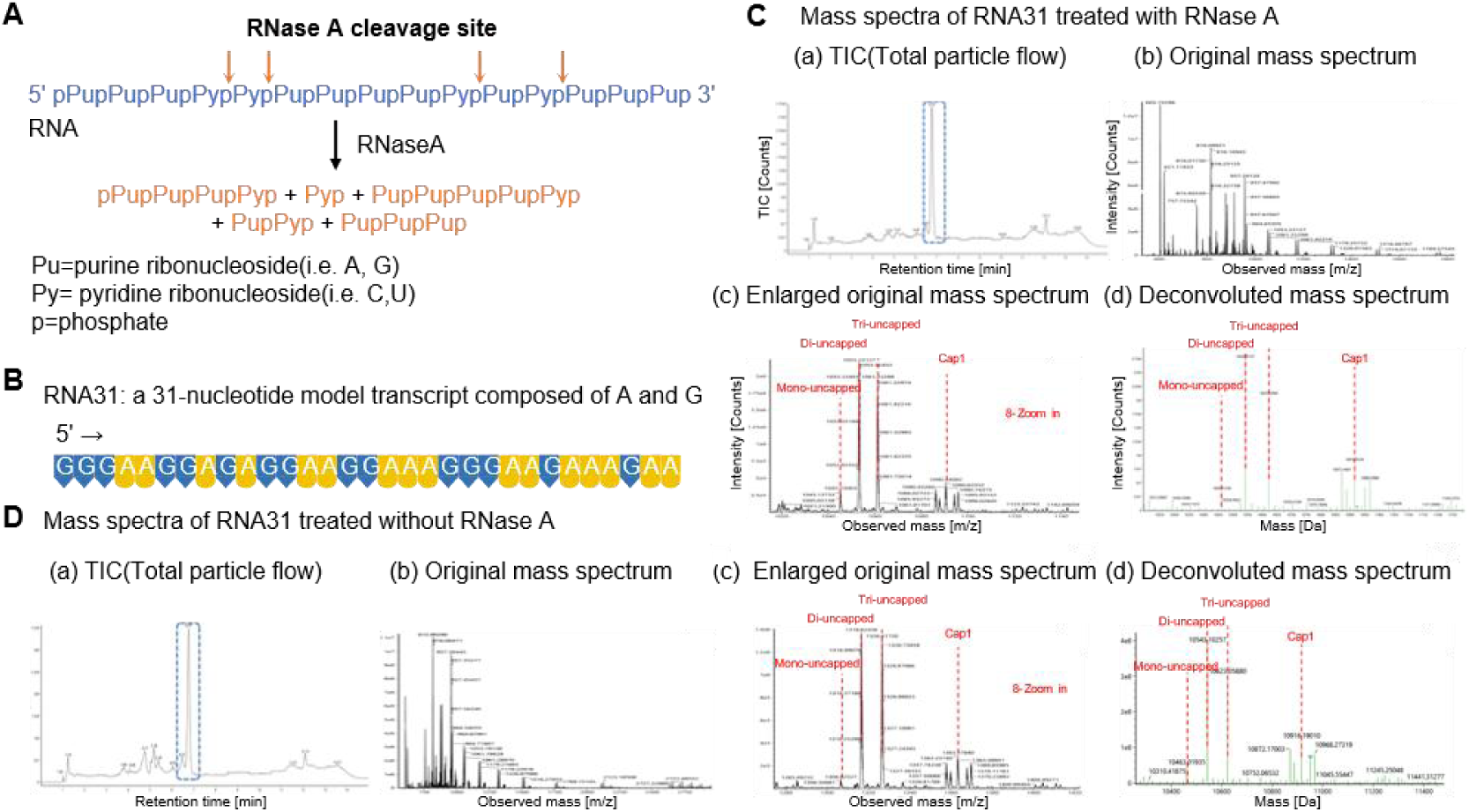
Specificity of RNase A cleavage and assessment of 5′ cap structural integrity. **(A)** Schematic mechanism of RNase A endoribonuclease activity, which selectively cleaves phosphodiester bonds 3’ of pyrimidine (Py: C, U) residues. (B) Primary sequence of a purine-only 31-nucleotide model transcript (RNA31) designed to verify 5’ cap stability under RNase A exposure. **(C, D)** LC-MS characterization of Cap1-capped RNA31 **(C)** following RNase A incubation and **(D)** without RNase A treatment, displaying total ion chromatograms (a), raw mass spectra (b), enlarged m/z regions (c), and deconvoluted mass spectra (d). Identical spectral profiles confirm that RNase A causes no non-specific cleavage or degradation of the 5’ Cap1 structure.

To confirm that RNase A digestion does not alter the capping status of capped RNA, we designed a 31-nucleotide model transcript, RNA31, composed exclusively of adenosine and guanosine residues (Fig. 2B, Supplementary Table S1, SEQ ID NO:3). RNA31 was prepared by IVT from a double-stranded DNA template generated by annealing complementary synthetic oligonucleotides (Supplementary Table S1, SEQ ID NO: 1 and 2, respectively). The transcription product was purified by reverse-phase HPLC and subsequently capped using a vaccinia capping enzyme system, followed by column purification to remove unreacted cap analogs and enzymes. Two sample sets were analyzed in parallel: an experimental group subjected to RNase A digestion under optimized high-salt conditions followed by spin-column purification, and an untreated control group processed identically but without enzymatic digestion. Both were analyzed by LC-MS/MS under identical instrumental conditions. Deconvoluted mass spectra of both groups revealed four principal RNA31 species corresponding to distinct 5’-terminal states: uncapped monophosphate (pRNA31, theoretical neutral mass 10,463.49 Da), uncapped diphosphate (ppRNA31, 10,543.49 Da), uncapped triphosphate (pppRNA31, 10,623.49 Da), and Cap1 (m^7^GpppGpA31, 10,916.49 Da) (Fig. 2C). Additional peaks shifted by +329 and +345 Da from the principal species were detected in both groups, consistent with non-covalent adenosine and guanosine adducts—a well-documented electrospray ionization (ESI) artifact in oligonucleotide mass spectrometry. Capping efficiency was calculated as the combined peak area of the Cap0 and Cap1 species divided by the total peak area of all detected RNA31 forms. In the RNase A-treated group, capping efficiency was determined to be 12.30% (Fig. 2C & Fig.S1B); in the untreated control, it was 12.76% (Fig. 2D & Fig.S1A). The 0.46 percentage-point difference falls within the expected analytical variability of the LC-MS/MS method (CV < 5%), confirming that RNase A treatment neither degrades the cap-bearing oligonucleotide nor alters the relative proportions of capped versus uncapped species. These results establish that RNase A digestion is fully compatible with downstream capping efficiency analysis.

### RNase A digestion enables label-free detection of the mRNA 5’ cap (Cap0/Cap1)

A functional IVT mRNA comprises, in the 5’→3’ direction, five structural elements: a 5’ cap structure, a 5’ UTR, an ORF, a 3’ UTR, and a poly(A) tail (Fig. 3A). The 5’ UTR is partitioned into two functionally distinct segments (Fig. 3A). Part A spans 9-27 nt and consists exclusively of adenosine and guanosine residues, terminating at its 3’ end with a cytidine or uridine nucleotide. This terminal pyrimidine serves as the sole RNase A cleavage site within the entire 5’-proximal region. Upon digestion, RNase A cleaves precisely at the Part A-Part B junction, releasing the 5’ cap-bearing fragment as a discrete analytical product. This fragment encompasses the cap structure (GpppN, Cap0, or Cap1) covalently attached to the Part A purine-rich oligonucleotide. The purine-only composition of Part A confers an additional functional benefit: the absence of internal pyrimidines ensures that no unintended cleavage occurs within the released cap fragment, preserving its integrity for subsequent mass spectrometric analysis. The poly(A) tail is engineered as a multi-block heteropolymeric structure composed of repeating AₘGₙ motifs, where each block contains m ≥ 30 adenosine residues interspersed by n = 1-4 guanosine spacer residues (Fig. 3A). This design exploits the same RNase A resistance principle for Part A. Because both adenosine and guanosine are purines, the poly(A) tail lacks pyrimidine residues and therefore contains no RNase A cleavage sites. Consequently, the poly(A) tail remains intact throughout the digestion reaction, irrespective of its total length (Fig. 3A).

**Fig. 3.**
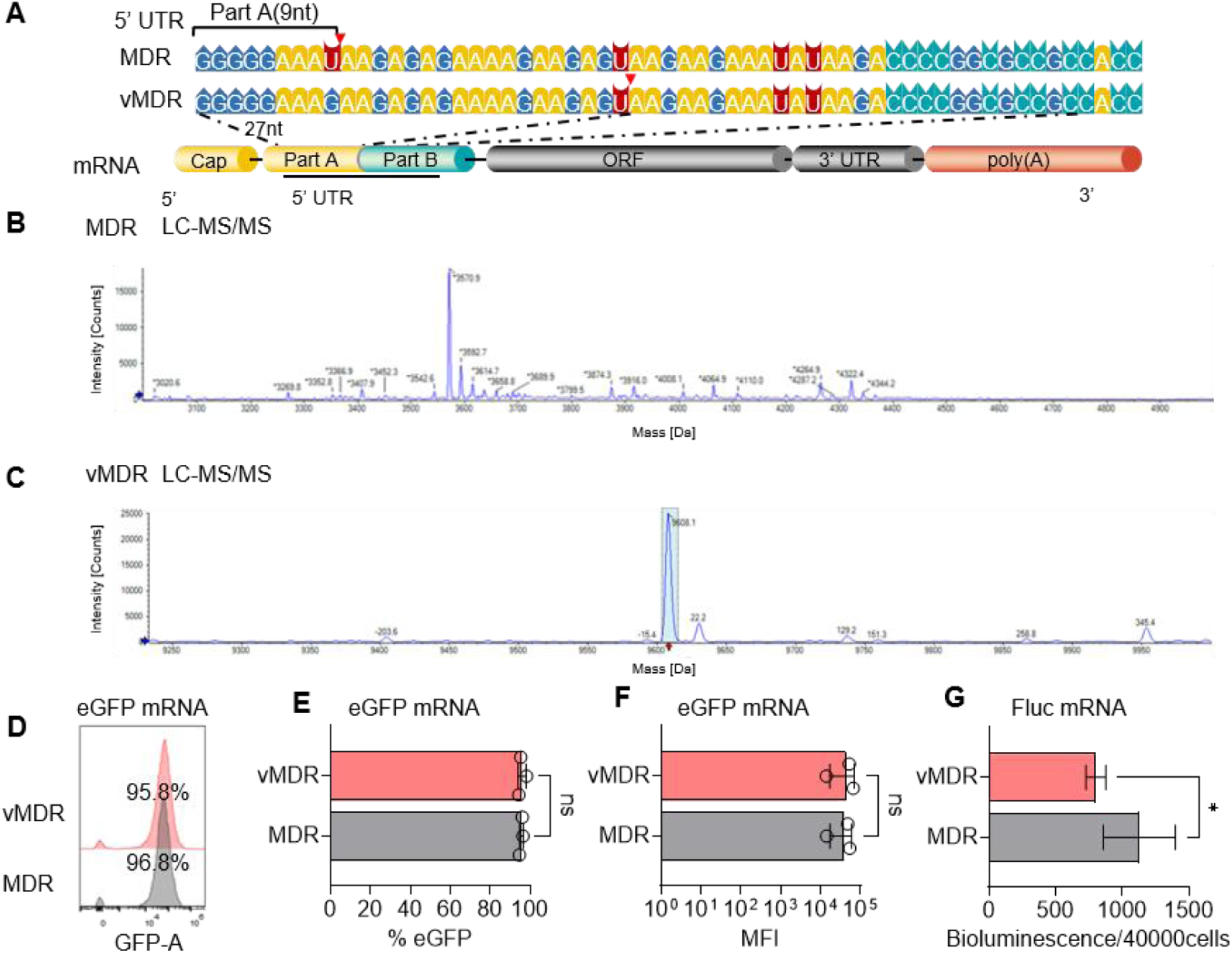
RNase A-directed 5’-UTR cleavage for capping efficiency assessment and translational characterization. **(A)** Schematic design of sequence-specific RNase A cleavage within MDR and modified vMDR 5’-UTRs, generating distinct 5’-terminal Part A fragments (9 nt and 27 nt, respectively). **(B, C)** LC-MS/MS mass spectra of cleaved 5’-terminal fragments from **(B)** MDR and **(C)** vMDR transcripts for capping efficiency quantification. **(D–F)** Flow cytometric analysis of eGFP reporter mRNA translation displaying **(D)** representative histograms, **(E)** percentage of eGFP-positive cells, and **(F)** mean fluorescence intensity (MFI). **(G)** Bioluminescence quantification of Fluc reporter mRNA translation (*P < 0.05; ns, not significant).

To assess whether the RNase A-based assay accommodates variable Part A lengths in the reporter mRNA, engineered 5’ UTR variants were designed by modifying the mRNA-1273 (Moderna-derived reference, MDR) 5’ UTR (Supplementary Table S1, SEQ ID NO: 18)^34^: the parental construct MDR (Part A = 9 nt) and its variant vMDR-eGFP (Part A = 27 nt), generated by a single U→G substitution at position 9 of the 5’ UTR that extends the purine-only tract from 9 to 27 nucleotides. Each variant was incorporated into enhanced green fluorescent protein (eGFP) reporter mRNA constructs and subjected to the one-step RNase A digestion workflow followed by LC-MS/MS analysis.

For MDR-eGFP (Part A = 9 nt), deconvoluted mass spectrometry revealed a predominant signal at 3570.9 Da, assigned to the Cap1-bearing 5’-terminal oligonucleotide, along with the expected sodium adduct peak at 3592.7 Da (Fig. 3B). Non-specific background signals were identified in the low-mass region but did not interfere with target peak integration. Capping efficiency was determined to be 97.6%. For vMDR-eGFP (Part A = 27 nt), the Cap1-bearing fragment was detected at 9608.1 Da, accompanied by the sodium adduct at 9630.4 Da (Δ +22.3 Da), consistent with Na⁺ adduction arising from sodium-containing mobile phase components (Fig. 3C). Capping efficiency was 98.2%.

These results demonstrate that the RNase A-based assay robustly quantifies capping efficiency across a broad range of Part A lengths (9–27 nt), with all tested constructs achieving capping efficiencies exceeding 97%.

A critical requirement for the analytical-compatible 5’ UTR design is that the engineered Part A segment must not diminish the translational performance of the mRNA. To verify this, we compared the protein expression of two eGFP/ Fluc reporter mRNAs. Both mRNAs were electroporated into 293T cells. eGFP expression was quantified by flow cytometry at 24 h post-electroporation, with mean fluorescence intensity (MFI) and the percentage of eGFP-positive cells serving as dual readouts. Both MDR-eGFP and vMDR-eGFP produced comparable eGFP expression profiles. The MFI values for the 9-nt and 27-nt Part A variants were 39309 ± 21823.41 and 44571 ± 27173.74, respectively (n = 3 biological replicates; P > 0.05, unpaired two-tailed t-test) (Fig. 3F). Similarly, the percentage of eGFP-positive cells did not differ significantly between the two constructs (96.45% vs. 96.05%; P > 0.05) (Fig. 3D-E). Importantly, Fluc expression was quantified at 24 h post-electroporation. The bioluminescence/40000cells values for the MDR-Fluc and vMDR-Fluc were 1127.50 ± 269.82 and 801.17 ± 74.08, respectively (n = 6 biological replicates; P = 0.0131) (Fig. 3G). These results confirm that extending the purine-rich Part A segment from 9 nt to 27 nt can reduce translational output in a reporter-dependent manner when the downstream Part B sequence and all other structural elements are held constant.

As anticipated, RNase A can be applied to mRNA samples for selective digestion of pyrimidine-containing regions without introducing measurement bias in capping efficiency, provided that the 5’ cap-bearing region is encoded within a purine-only segment. Further, we need to explore how to modulate Part A length without a translational penalty provides the flexibility to optimize the RNase A cleavage product size for different mass spectrometry platforms, ensuring broad applicability of the one-step dual-attribute assay across diverse mRNA therapeutic candidates.

### Rational engineering of the 5’ UTR enables RNase A-based detection of mRNA 5’ cap structures

Building on the mRNA-1273 (Moderna-derived reference, MDR) 5’ UTR, we rationally designed two additional UTR variants to evaluate the lower size boundary of Part A required for successful LC-MS/MS-based capping efficiency determination and to assess the impact of sequence modifications on mRNA translational output. The first variant, T1 (Fig. 4A; Supplementary Table S1, SEQ ID NO: 20), was designed with a Part A segment of 6 nt, below the shortest Part A length (9 nt) validated in the experiments described above. The second variant, T2 (Fig. 4A; Supplementary Table S1, SEQ ID NO: 21), was generated by systematically removing all 5’-proximal cytidine and uridine residues from T1, thereby extending the purine-only Part A tract to 23 nt and ensuring RNase A resistance across the entire cap-bearing region. Capping efficiency analysis of T1-eGFP mRNA (Part A = 6 nt) was performed using the established RNase A digestion and LC-MS/MS workflow. Theoretical neutral masses for the expected 5’-terminal oligonucleotide species were as follows: Cap1, 2551.3 Da; Cap0 (m⁷GpppN), 2537.3 Da; GpppN, 2272.3 Da. However, deconvoluted mass spectra (Fig. 4B) failed to reveal any detectable signal corresponding to these target masses, precluding capping efficiency calculation. In contrast, T2-eGFP mRNA (Part A = 23 nt) produced a clearly resolved deconvoluted mass spectrum (Fig. 4C). A predominant signal at 8243.55 Da was assigned to the Cap1-bearing fragment (theoretical neutral mass 8243.8 Da; mass deviation 30 ppm). An additional peak shifted by +337 Da was also detected, consistent with a non-covalent guanosine monophosphate adduct. Integration of peak areas across all detected 5’-terminal species yielded a capping efficiency of 99.5% for T2-eGFP mRNA. The stark contrast between T1 (undetectable) and T2 (99.5%) confirms that the Part A length threshold of ≥ 9 nt, established in our earlier systematic evaluation, represents a genuine analytical boundary rather than an artifact of sequence context. Furthermore, the successful detection of distinct Cap1 and adduct species at their expected masses validates the precision of RNase A cleavage at the designed pyrimidine (C/U) boundary. Had RNase A exhibited any non-specific endonuclease activity within the purine-rich Part A region, the resulting heterogeneous cleavage products would have produced a complex, unresolvable mass spectrum rather than the discrete species observed. This finding reinforces the conclusion that RNase A cleavage specificity under the optimized high-salt conditions (≥ 0.3 M NaCl) is sufficient to generate homogeneous analytical products.

**Fig. 4.**
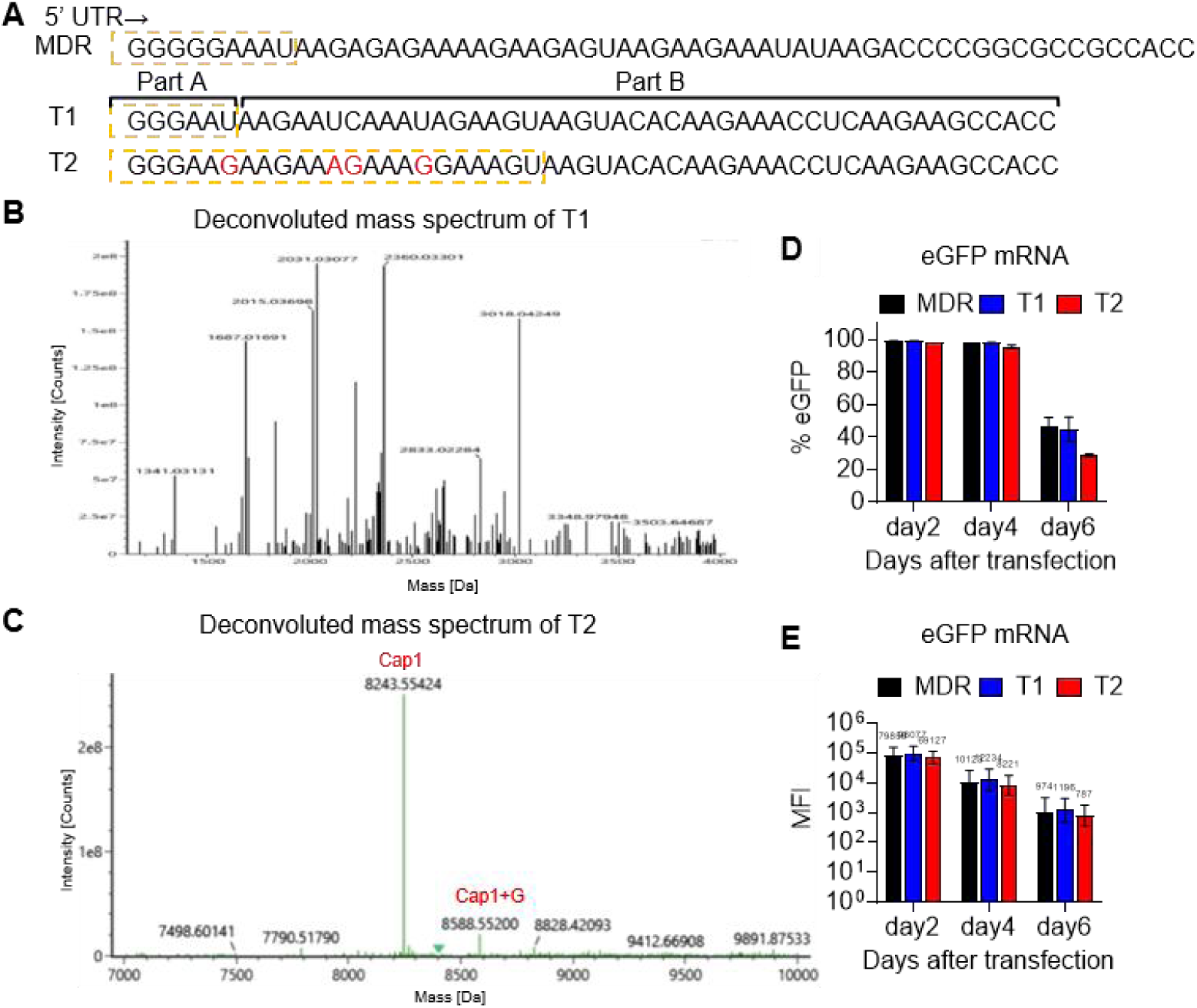
Rational design of synthetic 5’-UTRs for RNase A-based capping efficiency determination. **(A)** Sequence engineering of MDR-derived 5’-UTRs (T1 and T2). Pyrimidine-to-purine substitutions (red) in T2 eliminate premature cleavage sites, extending the 5’-terminal Part A fragment from 6 nt to 23 nt. **(B, C)** Deconvoluted mass spectra following RNase A digestion of **(B)** T1, which fails to yield identifiable capped fragments due to sub-optimal length, and **(C)** T2, enabling precise LC-MS resolution of Cap1 and Cap1+G species. **(D, E)** Time-dependent translational kinetics of eGFP reporter mRNAs carrying MDR, T1, or T2 5’-UTRs up to 6 days post-transfection, displaying **(D)** percentage of eGFP-positive cells and **(E)** mean fluorescence intensity (MFI).

To evaluate whether the sequence modifications introduced for analytical compatibility affect mRNA function, eGFP reporter constructs harboring the MDR, T1, or T2 5’ UTR were transfected into 293T cells, and eGFP expression was monitored at day2, day4, and day6 post-electroporation (Fig. 4D). The T1 construct, despite carrying the shortest Part A segment, produced eGFP levels comparable to those of the MDR reference across all three time points, indicating that the artificial T1 5’ UTR does not compromise translational efficiency relative to the benchmark design. In contrast, the T2 construct, which incorporates the C/U-to-purine substitutions necessary to extend Part A to 23 nt, exhibited a modest but reproducible reduction in eGFP expression relative to both MDR and T1.

Collectively, these results reveal that Part A segments of ≥ 9 nt are required for reliable LC-MS/MS detection, and while purine-only engineering of Part A is functionally tolerated, aggressive C/U removal may incur a translational penalty. These findings motivated subsequent engineering and screening of 5’ UTR variants designed to restore and further enhance translational output.

### Optimization of the 5’UTR for enhanced mRNA translation

UTR engineering represents a central strategy for optimizing the translational performance of IVT mRNA therapeutics. N^6^-methyladenosine (m^6^A) installed in the 5’ UTR has been reported to promote translation initiation through direct recruitment of eukaryotic initiation factor 3 (eIF3) and engagement of YTH-domain family reader proteins^35,36^. However, the precise mechanism and positional dependence of this effect remain debated, as recent structural and biochemical studies have questioned whether a single 5’ UTR m^6^A is sufficient to enhance translation initiation under homeostatic conditions^37^.

To empirically investigate the positional effect of m^6^A within our engineered 5’ UTR, three T2-derived variants were constructed by introducing a single m^6^A site at distinct positions within 5’ UTR: at the 5’-proximal end (T5), the central region (T4), and the 3’-distal end (T3) (Fig. 5A). Each variant was incorporated into an eGFP reporter mRNA construct and evaluated alongside the parental T2 construct lacking m^6^A.

**Fig. 5.**
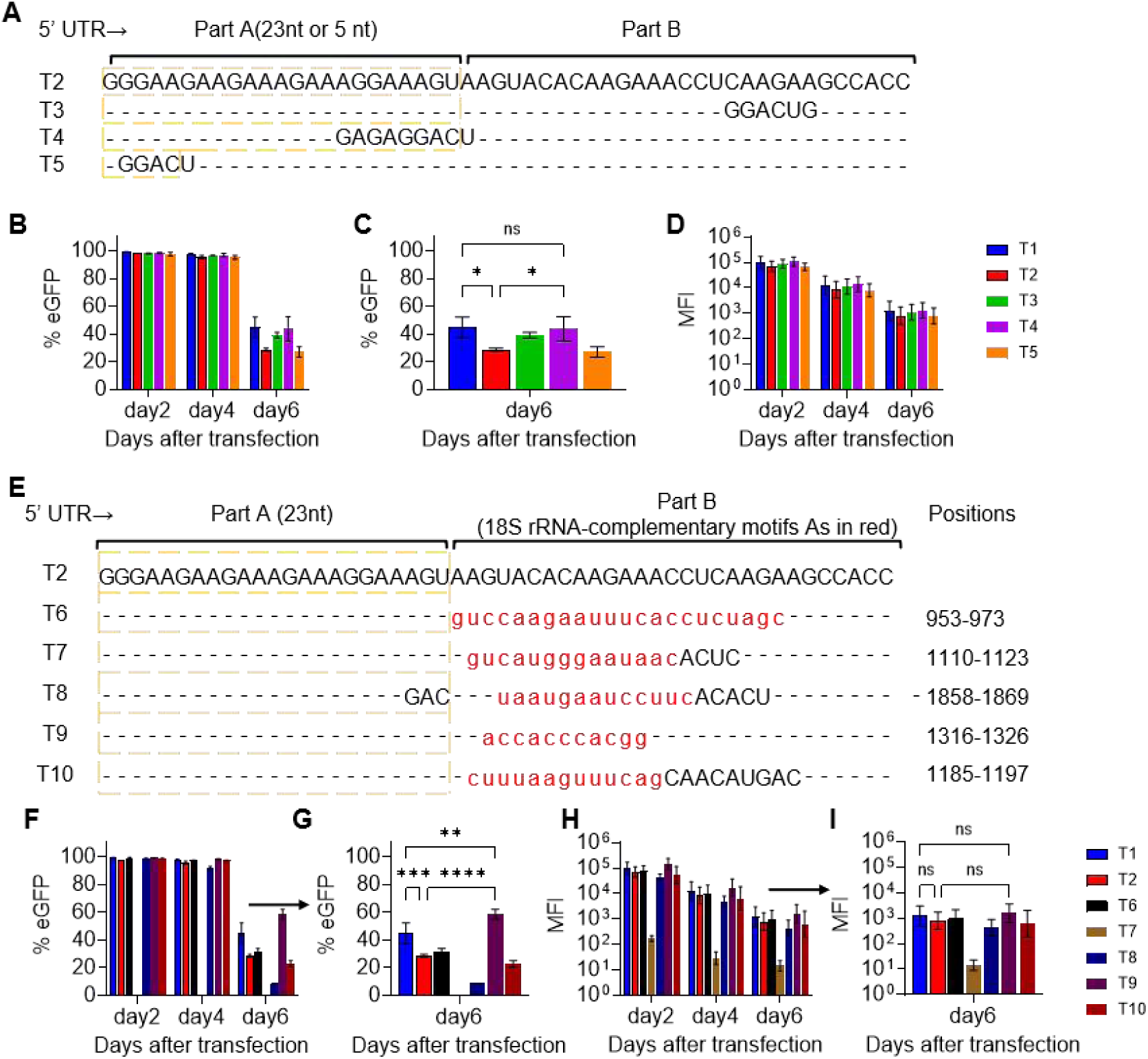
Enhancement of mRNA translational efficiency through engineered 5’-UTR motifs. **(A)** Sequence alignment of 5’-UTRs (T3–T5) containing m^6^A consensus motifs (GGACU) inserted at 3’-distal, central, and 5’-proximal regions. **(B–D)** Translational kinetics of m^6^A-modified eGFP transcripts (T1–T5) showing **(B, C)** percentage of eGFP-positive cells and **(D)** mean fluorescence intensity (MFI) over 6 days post-transfection. **(E)** Design of 5’-UTRs (T6–T10) incorporating 18S rRNA-complementary sequences. **(F–I)** Translational evaluation of 18S-targeted transcripts displaying **(F, G)** positive cell fractions and **(H, I)** MFI dynamics (*P < 0.05, P < 0.01, ***P < 0.001, ****P < 0.0001; ns, not significant).

Transfected 293T cells were monitored for eGFP expression at day2, day4, and day6 post-electroporation. At day 2 and day 4, all constructs achieved eGFP-positive cell rates exceeding 95%, precluding meaningful comparison (Fig. 5B). At day 6, however, a clear positional effect emerged: all three m^6^A-containing variants (T3, T4, T5) exhibited higher eGFP expression than the parental T2 construct, with the centrally positioned m^6^A variant (T4) producing the highest expression level (Fig. 5C). Mean fluorescence intensity (MFI) data across all groups were consistent with the positive-cell-rate trends (Fig. 5D). Notably, the T4 construct achieved expression levels comparable to those of the non-modified T1 construct (Part A = 6 nt, no C/U removal), suggesting that the translational penalty incurred by C/U removal in T2 can be at least partially recovered by strategic placement of m^6^A in the central region of the 5’ UTR.

Computational analyses of eukaryotic mRNA–ribosome interactions have identified evolutionarily conserved patterns of sequence complementarity between 5’ UTRs and 18S rRNA (NCBI Reference Sequence: NR_003286.4), suggesting that mRNA–rRNA base-pairing may modulate ribosome scanning and translation initiation efficiency^38,39^. Consistent with this model, we hypothesized that incorporating short sequences complementary to 18S rRNA into Part B of our engineered 5’ UTR could enhance ribosome recruitment during the scanning phase. Five distinct 18S rRNA-complementary motifs (each 11–21 nt) were individually inserted into Part B of the T2 5’ UTR backbone, generating variants T6–T10 (Fig. 5E). The specific complementary sequences targeting distinct regions of 18S rRNA are listed in Supplementary Table S1, SEQ ID NO: 43–47. All constructs were incorporated into eGFP reporter mRNA, prepared by in vitro transcription with capping and polyadenylation, and delivered into 293T cells by electroporation. eGFP expression was quantified by flow cytometry at day 2, day 4, and day 6 post-electroporation, with both the percentage of eGFP-positive cells and mean fluorescence intensity (MFI) serving as dual readouts.

Across all constructs, eGFP-positive cell rates exhibited a gradual decline over time, consistent with the transient nature of mRNA expression following electroporation (Fig. 5F). At day 2 and day 4, four of five 18S rRNA-complementary constructs (T6, T8, T9, T10) maintained positive-cell rates above 95%. However, variant T7, which harbors the complementary sequence 5’-GUCAUGGGAAUAAC-3’ (targeting positions 1110–1123 of human 18S rRNA), showed no detectable fluorescence signal at any time point, indicating complete translational failure or rapid mRNA degradation (Fig. 5F). At day 6, a clear differential emerged among the remaining constructs: T9, incorporating the motif 5’-ACCACCCACGG-3’ (targeting positions 1316–1326 of human 18S rRNA), achieved the highest positive-cell rate among all experimental groups and significantly outperformed both the T1 and T2 reference constructs (Fig. 5G). MFI values across all groups mirrored the positive-cell-rate trends (Fig. 5H).

These results demonstrate that the translational effect of 18S rRNA-complementary motifs is highly sequence-position-dependent: whereas the T9 motif substantially enhances sustained expression at day 2 to day 6, the T7 motif abolishes detectable protein output entirely. This biphasic behavior is consistent with the dual regulatory capacity of mRNA–rRNA complementarity previously reported in the literature, where specific complementarities can either enhance or inhibit translation depending on their structural context and the 18S rRNA binding site involved^38^.

### A probe-free RNase A assay for mRNA capping efficiency: equivalent accuracy to the RNase H probe method with a streamlined workflow

In current mRNA QC practice, capping efficiency and poly(A) tail length are typically assessed as separate analytical workflows. Capping efficiency is conventionally determined by a probe-based method, in which a custom-designed chimeric DNA–RNA probe is hybridized to the mRNA 5’ terminus, followed by RNase H cleavage of the RNA/DNA heteroduplex, streptavidin magnetic bead capture of the biotinylated probe-fragment complex, and LC-MS/MS analysis of the released 5’ cap-bearing oligonucleotide^21^ (Fig. 6A, upper left). Poly(A) tail length is independently measured by oligo(dT) magnetic bead capture of the mRNA 3’ end, RNase A digestion of the captured fragment, and LC-MS/MS characterization of the liberated poly(A) tail^14^ (Fig. 6A, upper right). These two sequential workflows require distinct sample preparations, customized probe synthesis, and magnetic bead–based enrichment steps, collectively consuming approximately 9 h (excluding the 4–6 days required for probe synthesis) and ¥256 per 100 μg of mRNA (Table 1). In contrast, the engineered 5’ UTR described herein enables a unified RNase A-based workflow that simultaneously generates both analytical targets from a single digestion reaction (Fig. 6A, lower). No sequence-specific probe is required, and purification is accomplished with a standard silica spin column rather than magnetic beads, substantially reducing both cost and hands-on time. To validate the RNase A one-step assay against the conventional approach, we selected the optimized T9 5’ UTR construct for side-by-side comparison. For capping efficiency, the RNase A method yielded a value of 96.7% (theoretical mass of the Cap1-bearing fragment: 8573 Da; Fig. 6B), while the RNase H probe method yielded 98.7% (theoretical mass of the probe-released fragment: 13,906 Da; Fig. 6C). Across three independent replicates, the two methods showed no statistically significant difference (Fig. 6D; paired two-tailed *t*-test, *P* > 0.05). Similarly, for poly(A) tail length determination, the RNase A one-step method (Fig. 6E) and the conventional magnetic bead–RNase A method (Fig. 6F) produced concordant results with no difference.

**Fig. 6.**
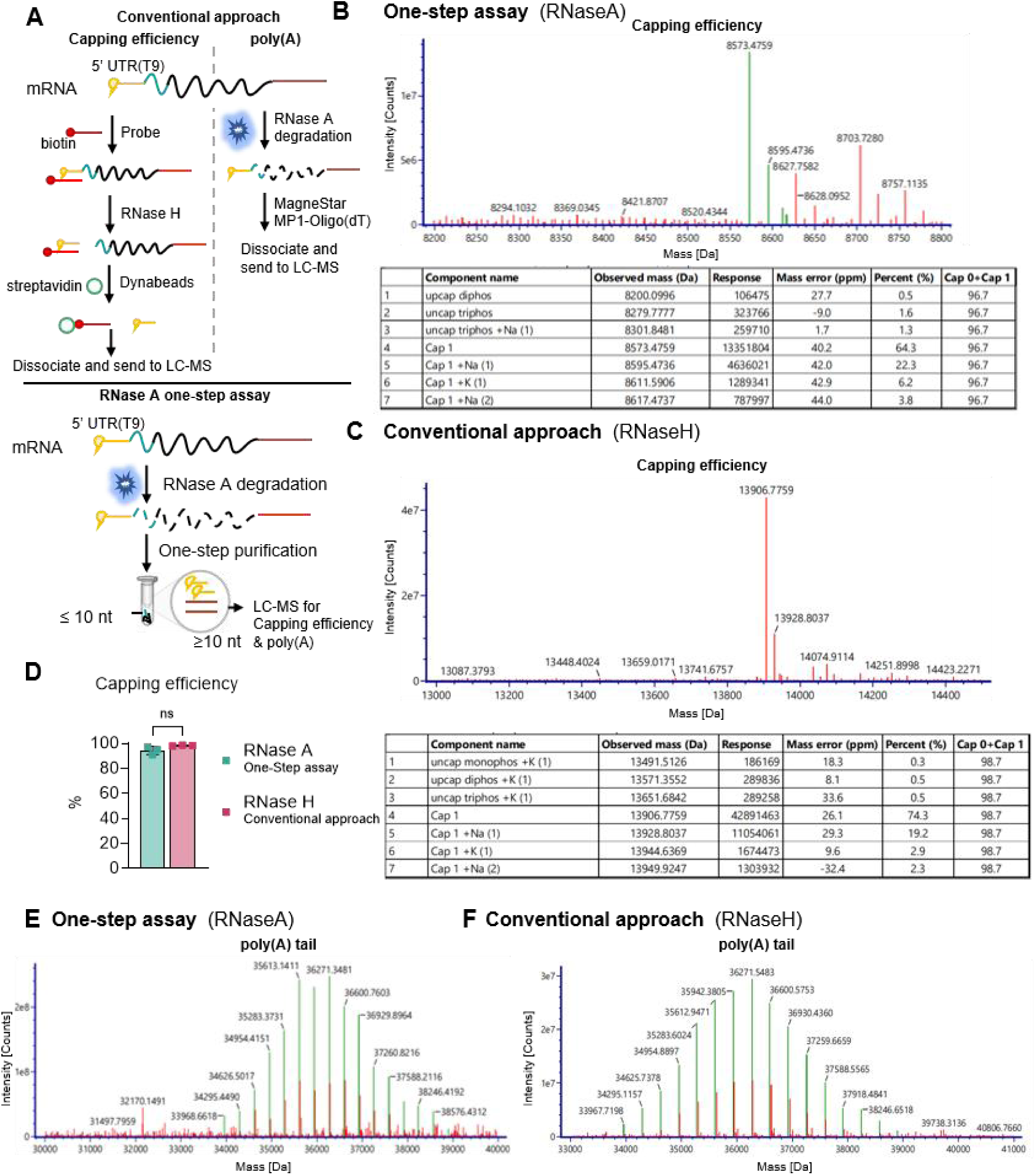
Methodological comparison of the RNase A one-step assay and conventional apporoach. **(A)** Schematics contrasting the multi-step probe-hybridization RNase H workflow against the streamlined RNase A one-step assay utilizing the engineered T9 5’-UTR transcript. **(B, C)** Representative deconvoluted LC-MS spectra and component tables for capping efficiency characterization obtained via **(B)** RNase A cleavage and **(C)** conventional RNase H mapping. **(D)** Cross-validation of total capping efficiency (% Cap0 + Cap1) between both analytical methods (*ns*, not significant). **(E, F)** Representative LC-MS spectra for poly(A) tail length determination characterization obtained via **(E)** RNase A cleavage and **(F)** conventional RNase.

**Table 1.** Economic and operational comparison between the RNase A one-step assay and the conventional approach for mRNA characterization. Reagent expenditure, processing duration, and workflow complexity per 100 μg of mRNA sample are evaluated. The RNase A one-step assay enables simultaneous assessment of capping efficiency and poly(A) tail length, reducing direct reagent costs by approximately 86% (∼¥35.43 vs ¥256.01) and cutting analytical turnaround time to one third (3 h vs. 9 h) relative to traditional probe-hybridization RNase H mapping.

| Method | Test items | Reagent | Price | Each 100 µg mRNA | Price/ ¥ | For 100 µg of mRNA |
| --- | --- | --- | --- | --- | --- | --- |
| RNase A one-step assay | Capping efficiency & poly(A) | RnaseA | 86¥/ ml | 5ul | 0.43 | The cost: approximately 35.43¥.<br>The processing time: approximately 3 hours. |
|  |  | Monarch® Spin RNA Cleanup Kits | 3500¥/ 100 preps | 1 preps | 35 |  |
| Conventional approach | Capping efficiency | RnaseH | 2079¥/ 50ul | 1 ul | 41.58 | The cost: approximately 256.01¥,<br>The processing time: approximately 9 hours(6 hours for Capping efficiency & 3 hours for poly(A) ). |
|  |  | Magnetic beads | 18000¥/ 10ml | 15 ul | 27 |  |
|  |  | Biotinylated DNA oligonucleotide probe | 8000¥/ 50OD | 4 ul | 160 |  |
|  | poly(A) | RnaseA | 86¥/ ml | 5ul | 0.43 |  |
|  |  | Magnetic beads | 18000¥/ 10ml | 15ul | 27 |  |

The cost and time advantages of the one-step assay are substantial. For 100 μg of mRNA, the RNase A method requires only RNase A (5 μL, ¥0.43) and one silica spin column (¥35), totaling ¥35.43 and approximately 3 h of processing time. The conventional approach requires custom probe synthesis (¥160), RNase H (¥41.58), and magnetic beads (¥27 for capping; ¥27.43 total for poly(A) analysis), amounting to ¥256.01 and approximately 9 h combined (excluding the 4–6 days required for probe synthesis). The RNase A method thus achieves a 7.2-fold reduction in per-sample cost and a 3-fold reduction in processing time (Table 1). Collectively, these data establish the RNase A one-step dual-attribute assay as a cost-effective, high-throughput-compatible alternative to the conventional two-workflow approach, eliminating the need for sequence-specific probe design and magnetic bead enrichment while delivering equivalent quantitative performance.

## Discussion

In this study, we have described an RNase A-based one-step dual-attribute assay that simultaneously quantifies mRNA capping efficiency and poly(A) tail length from a single enzymatic digestion, enabled by the rational engineering of the mRNA 5’ UTR. This approach addresses three longstanding limitations of current mRNA QC practice: (i) the requirement for separate, sequential analytical workflows for capping and poly(A) tail analysis; (ii) the sequence-dependent probe design for each target mRNA; and (iii) the substantial cost and time burden associated with custom probe synthesis and magnetic bead–based enrichment (Table 1).

The central analytical principle exploits the strict pyrimidine (C/U) 3’-specificity of RNase A: by encoding the 5’ cap-bearing region (Part A) and the poly(A) tail as pyrimidine-free, purine-only segments, both structures survive RNase A digestion intact and are co-purified by a single silica spin-column step for concurrent LC-MS/MS characterization. Our systematic evaluation established a Part A length threshold of ≥ 9 nt as the lower analytical boundary; the 6-nt Part A variant (T1) failed to produce detectable LC-MS/MS signals, likely attributable to the diminished ionization efficiency and chromatographic retention of very short oligonucleotides. Part A segments of 9–27 nt yielded robust, quantitative results across multiple mRNA constructs and reporter genes.

A head-to-head comparison with the conventional RNase H probe method, using the optimized T9 5’ UTR construct, demonstrated equivalent capping efficiency values (96.7% vs. 98.7%; P > 0.05, n = 3) and concordant poly(A) tail length determinations, while reducing per-sample cost by 7.2-fold and processing time by 3-fold. Notably, the RNase A assay eliminates the need for sequence-specific probe synthesis. This represents a critical advantage for high-throughput in-process control, where rapid turnaround across multiple mRNA candidates is essential.

This concept of simultaneous dual-CQA analysis was independently reported by Wang *et al*., who employed 10-23-type DNAzymes to generate both 5’ and 3’ end fragments in a single reaction^22^. The two strategies differ fundamentally in their design philosophy: the DNAzyme approach requires custom-designed catalysts for each mRNA sequence, whereas the present method leverages the intrinsic, sequence-independent cleavage specificity of RNase A, conditioned only on the presence of a pyrimidine-free Part A segment in the 5’ UTR. This distinction confers a practical advantage: once a pyrimidine-free Part A is incorporated into the mRNA template, no additional reagent design or synthesis is required, making the method readily generalizable across diverse mRNA therapeutic candidates.

A key design consideration revealed by our 5’UTR screening is the potential trade-off between analytical compatibility and translational performance. The purine-only Part A architecture is functionally neutral at lengths of 9-27 nt, consistent with the established permissiveness of unstructured, purine-rich 5’ UTR segments for cap-dependent translation^40^. However, comprehensive C/U removal extending Part A to 23 nt (T2) incurred a measurable translational penalty in 293T cells. This penalty was recoverable through two complementary strategies: (i) strategic placement of m^6^A in the central region of 5’UTR, which restored translational output to levels comparable with the unmodified T1 construct, consistent with the proposed role of 5’ UTR m^6^A in facilitating translation initiation complex assembly; and (ii) incorporation of 18S rRNA-complementary motifs into Part B, with variant T9 achieving the highest sustained expression at day6. The five tested complementary motifs exhibited divergent effects on translation. Notably, motif T9 enhanced protein expression, whereas motif T7 abolished detectable translational output. These contrasting outcomes are consistent with the dual regulatory capacity of mRNA–rRNA complementarity reported previously^38,39^ and underscore the necessity of empirical screening for each candidate motif.

In summary, the RNase A one-step dual-attribute assay provides a streamlined, cost-effective, and sequence-independent analytical platform for concurrent monitoring of two critical quality attributes of IVT mRNA. The engineered UTR architecture simultaneously supports efficient translation and analytical accessibility, establishing a design paradigm in which sequence elements serve dual functional and diagnostic roles within the mRNA therapeutic construct.

## Acknowledgements

### Funding

This study was supported by Shanghai Science and Technology Innovation Action Plan, Cell and Gene Therapy Project, Science and Technology Commission of Shanghai Municipality (23J21900500) awarded to P.J.Z. as the principal investigator.

## Author contributions

Y.F. developed the concept with input from P.J.Z. and Q.J.Q., Y.F. performed cell-based experiments, curated and interpreted all datasets, and wrote the manuscript. M.Q.S. constructed the template vectors for molecular experiments. T.L. designed and screened 5’ UTR sequences for high-efficiency translational expression. Y.W.L. prepared mRNA samples and performed capping efficiency analyses. X.D.W. conducted cell-based assays, prepared mRNA samples, performed capping efficiency measurements, and organized raw datasets. C.C.W. constructed template vectors, established the IVT system, and performed mRNA sample preparation and capping efficiency detection. J.C.Y. prepared mRNA template plasmids and IVT mRNA. P.J.Z. conceived and oversaw the overall study design, developed the rapid capping efficiency detection strategy, revised the manuscript, and served as the principal investigator of the funding project. Q.J.Q. provided funding acquisition and served as the academic supervisor at Shanghai University. All authors reviewed and approved the final manuscript.

We would like to thank Linxiao Du, Xiangxiang Bao for Biophysical characterization of capping efficiency and poly(A) with LC-MS.

